# Protein-protein interfaces as druggable targets: A common motif of the pyridoxal-5’-phosphate-dependent enzymes to receive the coenzyme from its producers

**DOI:** 10.1101/2023.02.13.528268

**Authors:** Vasily A. Aleshin, Victoria I. Bunik

**Affiliations:** Department of Biokinetics, A. N. Belozersky Institute of Physicochemical Biology, Lomonosov Moscow State University, 119234 Moscow, Russia; (V.A.A.); (V.I.B.); Department of Biochemistry, Sechenov University, 119048 Moscow, Russia; Faculty of Bioengineering and Bioinformatics, Lomonosov Moscow State University, 119234 Moscow, Russia

**Keywords:** CBS, PdxK, PNPO, PLPBP, pyridoxal-5’-phosphate-dependent enzyme, TAT, vitamin B6 metabolism, amino acid metabolism, 1-carbon metabolism

## Abstract

Pyridoxal-5’-phosphate (PLP), a phosphorylated form of vitamin B6, acts as a coenzyme for numerous reactions, including those changed in cancer and/or associated with the disease prognosis. Since highly reactive PLP may modify cellular proteins, it is hypothesized to be directly transferred from its donors to acceptors. Our goal is to validate the hypothesis by finding common motif(s) in a multitude of the PLP-dependent enzymes for binding the limited number of the PLP donors, namely pyridoxal kinase (PdxK), pyridox(am)in-5’-phosphate oxidase (PNPO) and the PLP-binding protein (PLPBP). Experimentally confirmed interactions between the PLP donors and acceptors reveal that PdxK and PNPO interact with the PLP acceptors of folds I and II, while PLPBP – with those of folds III and V. Aligning the sequences and 3D structures of the identified interactors of PdxK and PNPO, we have found a common motif in the PLP-dependent enzymes of folds I and II. The motif extends from the enzyme surface to the neighborhood of the PLP binding site, represented by an exposed alfa-helix, a partially buried beta-strand and residual loops. Pathogenicity of mutations in human PLP-dependent enzymes within or in the vicinity of the motif, but outside of the active sites, supports functional significance of the motif that may provide an interface for the direct transfer of PLP from the sites of its synthesis to those of the coenzyme binding. The enzyme-specific amino acid residues of the common motif may be useful to develop selective inhibitors blocking PLP delivery to the PLP-dependent enzymes critical for proliferation of malignant cells.

## INTRODUCTION

The enzymes using pyridoxal-5’-phosphate (PLP) as the coenzyme encompass more than 300 distinct catalytic functions, belonging to five of the seven existing EC classes [1]. Many of these reactions, particularly those involving amino acid metabolism, are critical for not only metabolic switches underlying malformations, but also the elimination of malignant cells. For instance, PLP-dependent transaminases drive T cells activation and differentiation, required for anticancer responses [2]. Down-regulation of the PLP-dependent tyrosine aminotransferase (TAT) encoded by *TAT* gene, supports pathogenicity of hepatocellular carcinoma [3], with the low expression of the tyrosine catabolic enzymes predicting poor outcome [4]. Different cancer types exhibit changes in the expression of the PLP-dependent cystathionine β-synthase (CBS), associated with poor prognosis, in accord with the enzyme essential role in one-carbon metabolism including the methylation and transsulfurylation reactions [5, 6].

PLP is the major component of the B6 vitamers pool, estimated to be in the micromolar concentration range in serum and red blood cells [7, 8]. PLP is hydrolyzed to pyridoxal (PL), either non-enzymatically or by numerous phosphatases, including PLP-specific phosphatase (PLPP, or chronopin) that is under the control of hypoxia-inducible transcription factor HIF1 regulating metabolism of T cels [2]. Among the PLP donor proteins, PLP-binding protein (PLPBP, gene *PLPBP*) covalently binds PLP by formation of a Schiff base between PLP aldehyde group and PLPBP lysine residue [9], while pyridoxal kinase (PdxK, gene *PDXK*) and pyridox(am)in-5’-phosphate oxidase (PNPO, gene *PNPO*) are the two PLP-producing enzymes [10, 11].

Not only the PLP-dependent reactions, but also the metabolism of vitamin B6 are changed in cancer. Markers of the inflammation-induced degradation of the PLP precursor PL correlate with cancer incidence, especially for the lung cancer that is significantly contributed by inflammation [12]. Formation of PLP from PL by PdxK, activities of PLP-dependent ornitine decarboxylase (*ODC1* gene) and mitochondrial aspartate aminotransferase (*GOT2* gene) are critical for specific metabolic changes supporting proliferation of myeloid leukemia cells [13]. Levels of PdxK protein increase in the lung cancer, in contrast to the PdxK mRNA levels, and the increase correlates with good prognosis [14]. Overall, analysis of the changes in metabolism of vitamin B6 in different cancer cells reveals a very complex relationship between the changes and disease progression. Although B6 deficiency may be deleterious for cancer cells, it also promotes malignant transformation of normal cells, stimulating DNA damage and compromising immune response [15]. Pharmacological regulation of specific PLP-dependent reactions may provide new tools to selectively fight various cancer types through their essential dependence on some of these reactions. However, current inhibition of the PLP-dependent processes is based on chemical reactivity of PLP and lacks the a desired enzyme specificity.

Both PLP an its catalytically inactive precursor PL are reactive aldehydes, that can modify protein lysine residues or amino groups of low molecular weight compounds, such as a widely used inhibitor of PLP-dependent transaminases, aminoxyacetic acid. These reactions explain toxicity of PLP and PL in excessive concentrations. For instance, at 0.5 mM, these vitamers are much more effective in killing varous cancer cells, than pyridoxamine or pyridoxin [16]. In view of the PLP reactivity, it has been hypothesized that PLP is transferred directly from its donors to the enzymes which use PLP as the coenzyme (PLP acceptors) [17–19]. However, the structurally different proteins employing PLP as their coenzyme, encompass up to seven structural folds without significant homology [1, 20]. Given the three PLP donor proteins, i.e. PdxK, PNPO and PLPBP, the multitude and structural variety of the PLP acceptor proteins imply that there are subsets of the acceptors possessing common structural motifs for binding PdxK, and/or PNPO, and/or PLPBP. The goal of this work is to assess, if these common structural motif(s) implied by the hypothesis about the direct transfer of PLP from its donors to acceptors, may be found in the structurally different PLP acceptors. Based on the database and literature search, we identify experimentally confirmed physical interactions between the PLP acceptors and donors, further analyzing the non-homologous PLP acceptor proteins for the local similarity of their sequences and 3D structures. The common motif is thus revealed in the PLP-dependent enzymes of the structurally different folds I and II, that may provide an interface for the PLP acceptors binding of PdxK or PNPO. The interface may be employed for PLP channeling from the donors to acceptors. The functional significance of the found motif is supported by pathogenic mutations within the motif or its neighborhood outside the active sites, known for monogenic diseases associated with human variants of the PLP-dependent cystathionine β-synthase (CBS) or tyrosine aminotransferase (TAT) [21, 22]. Since changed expression of CBS and TAT is known to be associated with malignant transformation and poor prognosis, pharmacological regulation of the protein-protein interactions supporting the PLP transfer to these enzymes, may represent a novel strategy to develop the enzyme-specific drugs. Such strategy may use specific elements of the identified common motif to block the interface presumably involved in the PLP transfer.

## MATERIALS AND METHODS

### Bioinformatics motif search

The sequence-based search for conserved motifs in the PLP-dependent enzymes uses GLAM2, version 5.4.1 [23], available at the online server (https://meme-suite.org, accessed 25 Sept. 2022). Given selected sequences as input, this algorithm finds the motifs through gapped local alignments. The input sequences have been identified by our search of the published data on the experimentally identified interaction partners of the PLP donors. The recommended parameters of the algorithm are used, but the motif is forced to be found in all sequences.

### Structural visualization and alignment

PyMOL v1.7 (PyMOL Molecular Graphics System, Schrödinger, LLC.) is used for the visualization of proteins as described in the corresponding figure legends. The proteins are represented as cartoon models, with different chains shown in different colors of gray. The standard color code is used for ions or ligands. The structures are downloaded from the Protein Data Bank. Protein-BLAST [24] is used to identify the homologs of proteins available in the database, if necessary.

To retrieve the sequences of mammalian fold II proteins corresponding to the motif of diaminopropionate ammonia-lyase (dpaL) protein, the structural alignment of the proteins by PyMOL is used. The sequences corresponding to the motif of dpaL are further used to prepare the logo with the help of GLAM2.

### Multiple sequence alignment

Qlustal Omega multiple sequence alignment [25] is applied to align the sequences of homologous proteins (total identity > 30%).

## RESULTS

### Interactions of PNPO, PdxK and PLPBP with PLP-dependent enzymes

Results of our search through publications and the open-access databases (BioGrid [26], IntAct [27], String [28]) on the confirmed physical interactions between the PLP-dependent enzymes, i.e. PLP acceptors, and the three known PLP donor proteins are shown in table. Nine of the twelve interactions refer to the PLP-dependent enzymes of the fold I, that is the most abundant fold type including more than 75% of the PLP-dependent proteins [1]. However, also the members of less abundant PLP-binding proteins of folds II, III, and V, are confirmed interactors of the PLP donors (table).

**Table.** Experimentally confirmed interactions of the PLP-dependent enzymes with the PLP-donating proteins.

| PLP donor | No | PLP-dependent enzyme | Organism, method | Fold | Refs |
| --- | --- | --- | --- | --- | --- |
| PdxK | 1 | 4-aminobutyrate aminotransferase, *ABAT*, P80404, EC: 2.6.1.19 | human, co-fractionation and mass-spectrometry | I | [29] |
|  | 2 | ornithine aminotransferase, *OAT*, P04181, EC: 2.6.1.13 | human, co-fractionation and mass-spectrometry | I | [29] |
|  | 3 | aspartate aminotransferase, *GOT1*, W5PS88, EC: 2.6.1.1 | sheep, emission anisotropy and affinity chromatography | I | [30] |
|  | 4 | alanine aminotransferase, *GPT*, A0A287BMB4, EC: 2.6.1.2 | pig, fluorescence polarization and surface plasmon resonance | I | [19] |
|  | 5 | glutamate decarboxylase, *GAD1*, P48319, EC: 4.1.1.15 | pig, fluorescence polarization and surface plasmon resonance | I | [19] |
|  | 6 | serine hydroxymethyltransferase, *glyA*, Q7SIB6, EC: 2.1.2.1 | *G. stearothermophilus*, surface plasmon resonance | I | [31] |
|  | 7 | diaminopropionate ammonia-lyase, *dpaL*, P40817, EC: 4.3.1.15 | *S. typhimurium*, surface plasmon resonance | II | [31] |
| PNPO | 8 | serine hydroxymethyltransferase, *SHMT1*, P07511, EC: 2.1.2.1 | rabbit, fluorescence polarization and kinetics | I | [32] |
|  | 9 | aspartate aminotransferase, *aspC*, P00509, EC: 2.6.1.1 | *E. coli*, fluorescence polarization and kinetics | I | [32] |
|  | 10 | L-threonine aldolase, *ltaE*, P75823, EC: 4.1.2.48 | *E. coli*, fluorescence polarization and kinetics | I | [32] |
| PLPBP | 11 | glycogen phosphorylase, *PYGB*, P11216, EC: 2.4.1.1 | human, chemical cross-linking and MS | V | [33] |
|  | 12 | pyridoxal phosphate homeostasis protein, *PLPBP*, O94903, | human, chemical cross-linking and MS | III | [33] |

\*PLP-dependent enzymes, are given together with the names of their genes and Uniprot identifiers.

The identified interactions reveal that proteins of fold I bind PdxK, PNPO or both. On the other hand, the most studied PLP donor, i.e. PdxK, with the highest number of protein partners may bind the PLP-dependent enzymes of the structurally different folds I and II. Finally, the two known interactors of PLPBP, including the PLPBP homo-oligomerization, belong to the different structural folds III and V. PLPBP and its bacterial orthologs show high structural similarity to N-terminal domain of some PLP-dependent enzymes, such as bacterial alanine racemase and eukaryotic ornithine decarxylase, but no enzymatic activity has been found for PLPBP [33, 34].Thus, the published data on the confirmed interactions between the PLP donors and acceptors indicate that (1) PdxK may channel their product PLP to the PLP acceptor proteins of folds I and II; (2) PdxK and PNPO may have common partners among the PLP acceptors of fold I; (3) PLPBP partners with the PLP acceptor proteins of rare structural folds III and V.

The sequence-based search for local similarity using GLAM2 (version 5.4.1) [23] has identified a motif, common to all the PLP-dependent interactors of PdxK (table, No 1-7), shown in fig. 1a. In good accordance to the existence of the common partners of PdxK and PNPO, such as serine hydroxymethyltransferase (SHMT) and aspartate aminotransferase (GOT) (table), a similar motif has been found in the search using the sequences interacting with both the PNPO and PdxK (fig. 1b). In this case, the motif becomes five residues shorter from its N-terminus and two residues longer on its C-terminus (fig. 1b *vs* 1a). That is, pooling the PdxK and PNPO interactors together does not change the major part of the common motif, slightly modifying only the terminal parts of the motif.

**Fig. 1.**
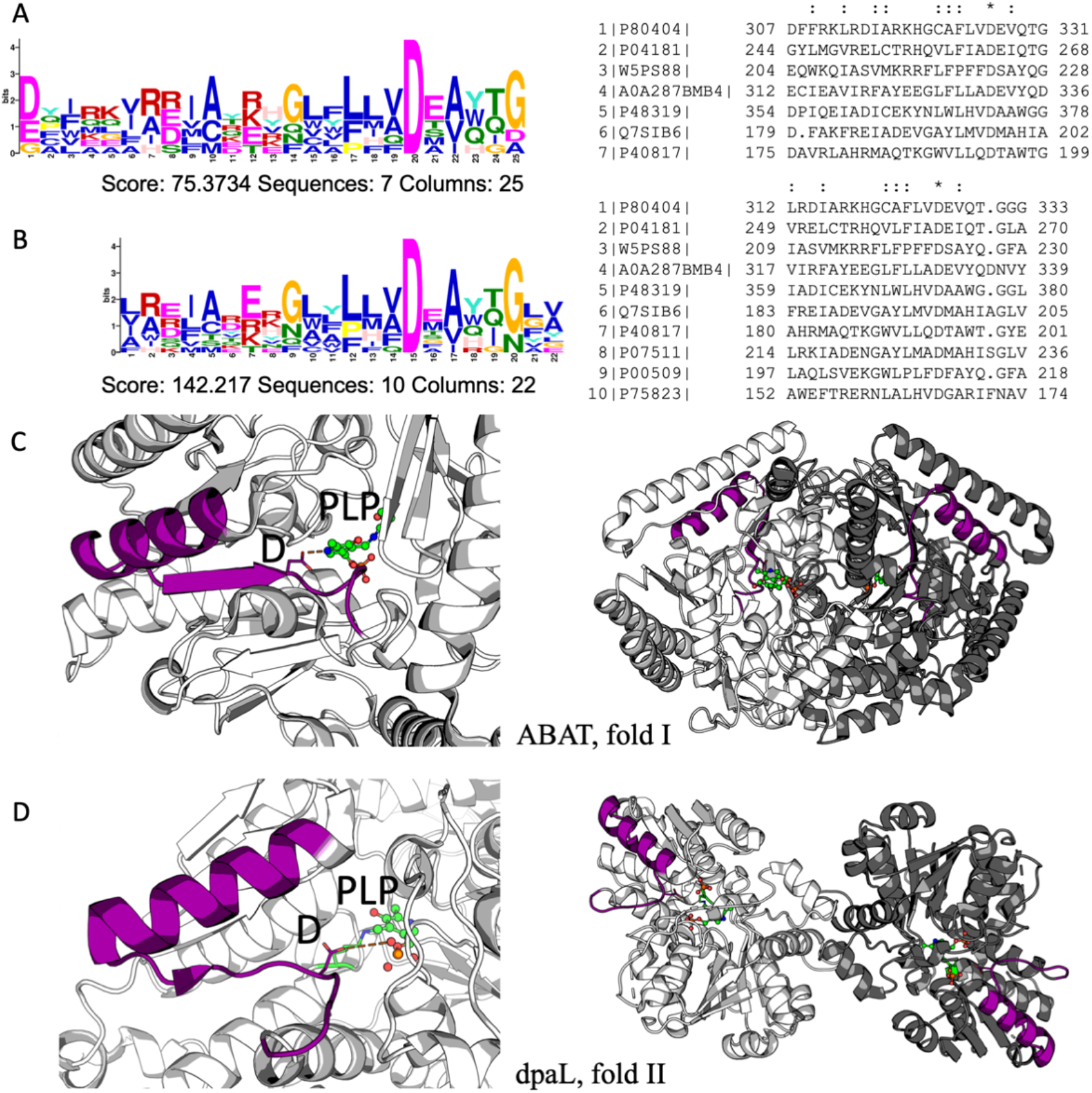
A common GLAM2-identified motif of the PLP-binding enzymes that are shown to interact with PLP donors PdxK and PNPO. (a) - The logo and sequences of the motif, found in the PdxK-binding mammalian and bacterial enzymes. (b) – The logo and sequences of the motif, found in the PdxK- and PNPO-binding mammalian and bacterial enzymes. (c) – A closer view of the motif (left) and its localization in the dimeric structure (right) of GABA transaminase (ABAT, fold I, structure: 4ZSW). (d) – A closer view of the motif (left) and its localization in the dimeric structure (right) of diaminopropionate ammonia-lyase (dpaL, fold II, structure: 5YGR). Numbers of the sequences in (a) and (b) are identical to those in the table.

Comparison of fig. 1c and 1d reveals that the found motif, common to all the partners of PdxK and PNPO, has a similar secondary structure in the PLP-dependent enzymes of different structural folds I and II. The motif is represented by an alpha-helix, a beta-strand and the interconnecting and C-terminal loop regions. The motif within the structures of all the PdxK and PNPO interactors listed in table, is visualized in Supplementary fig. S1. Remarkably, the revealed motif has been earlier identified as the structural signature of the fold I enzymes, with the beta-strand end including a conserved aspartate residue known to interact with PLP [20, 35]. Our work extends this common motif to the fold II enzymes, shown to interact with the same PLP donors as fold I enzymes (table, fig. 1). Importantly, the helix and loops of the motif are exposed to the surrounding medium, also in the enzyme dimers (fig. 1c,d, Supplementary fig. S1), pointing to spatial availability of the common motif to the heterologous protein-protein interactions.

Further, we have elaborated procedures to identify the motif in the PLP-dependent enzymes of the fold I and II, whose physical interactions with PLP producers are not known. As mentioned above, in all the enzymes of the structural fold I, the motif has been identified earlier as a common structural element neighboring the conserved PLP-binding aspartate residue of the active site [35]. Addition of the sequences of the fold I enzymes tyrosine transaminase (P17735) and kynurenine transaminase 1 (Q16773) to those of the 10 known interactors of PdxK and PNPO shown in table, followed by repeated GLAM2 search, finds the motif in the two unknown interactors, improving the GLAM2 score (Supplementary fig. S2).

Identification of the common motif of the PdxK/PNPO interactors among the enzymes of structural fold II is more challenging. Bacterial diaminopropionate ammonia-lyase (dpaL), where we have identified this motif (fig. 1d), has no orthologs in mammals. Hence, known human PLP-dependent enzymes of the structural fold II, such as serine racemase (SRR, Q9GZT4), serine dehydratase-like (SDSL, Q96GA7) and cystathionine β-synthase (CBS, P35520) (fig. 2) have been chosen for identification of the common motif found in diaminopropionate ammonia-lyase. The sequences of human enzymes have been added to those shown in fig. 1, separately or all together. Performing GLAM2 for these sets of the sequences of mammalian fold II proteins decreases the overall score, with the motif thus identified demonstrating no specific secondary structure. The poor performance of the sequence alignment agrees with the fact that the fold II proteins may possess very different protein sequences. For instance, CBS has a sequence extension for its heme binding, not employed by SRR or SDSL (fig. 2a). Besides, the aspartate residue in the motif of dpaL protein, conserved in the fold I proteins, is no more conserved in the fold II proteins. As seen from the alignment and corresponding logo in fig. 2b, this residue may be substituted by other residues, exemplified by the glutamine or proline residues in the shown human fold II proteins. Therefore, to identify the common structural motif in such structurally different proteins, structural alignment of dpaL protein and human enzymes of fold II is required. Using this approach, the motif similar to that of bacterial dpaL protein, is found in mammalian SRR, serine dehydratase-like and CBS (fig. 2a). Remarkably, also in the fold II enzymes, the C-terminal end of the common motif is vicinal to PLP (fig. 2a), similar to the fold I proteins where the conserved aspartate residue of the C-terminal end of the motif interacts with the PLP nitrogen atom (fig. 1). In SRR, also the N-terminal part of the motif contributes to the active site. That is, the amino acid residues at the start (R135) and the end (H152-P153-N154 loop) of the SRR motif are known to be involved in binding of SRR substrate and PLP-imine stabilization, correspondingly [36]. Participation of the terminal parts of the common motif in the active site formation supports the role of the motif in the delivery of PLP to the acceptor active sites from the PLP donors.

**fig- 2.**
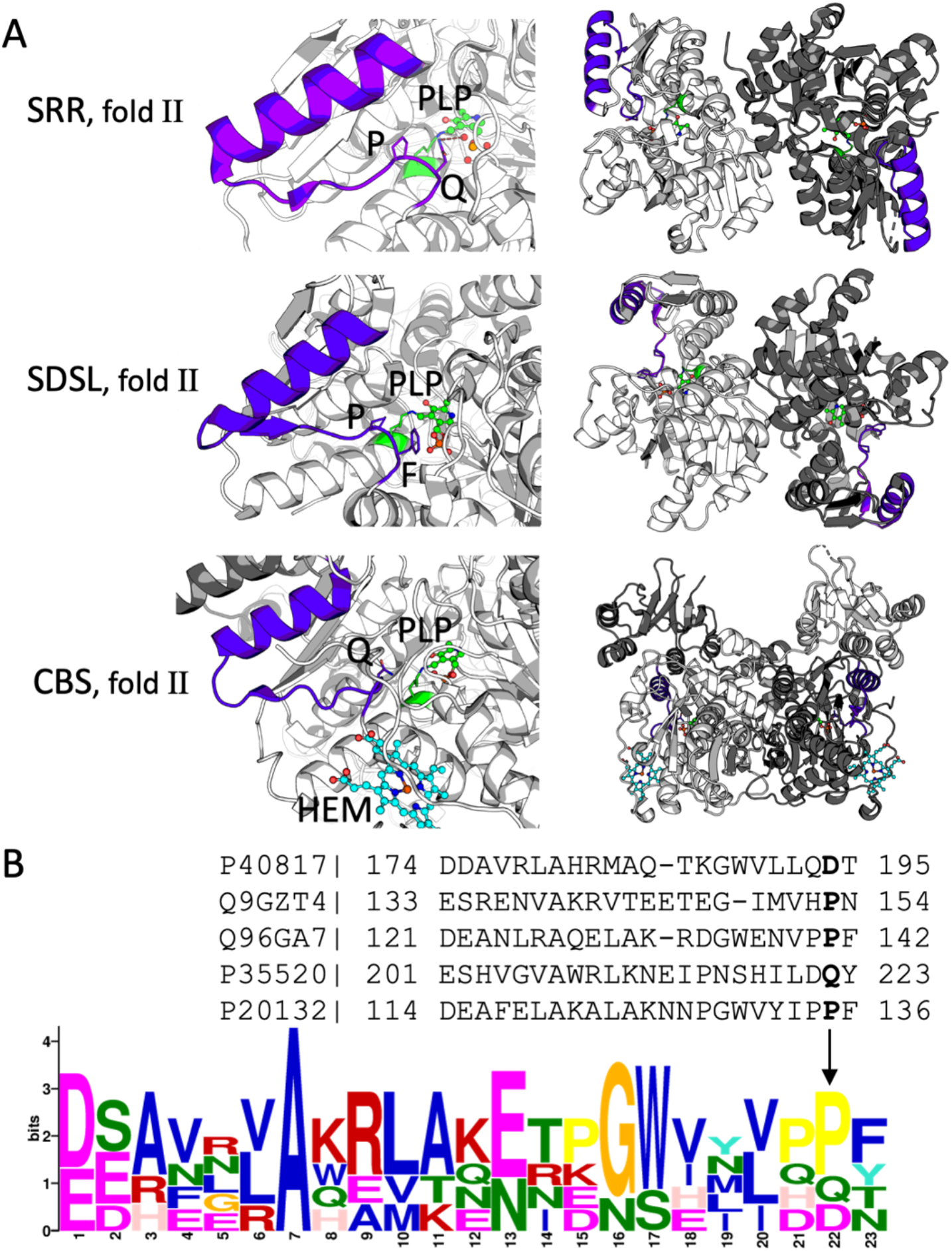
Identification of the structural motif, common for the PLP-dependent enzymes, in the mammalian fold II proteins, using the motif found in the experimentally confirmed PdxK interactor of fold II, dpaL protein. (A) - At the left, the common motif identified in the mammalian fold II proteins, is shown in blue, corresponding to the purple motif in dpaL protein, shown in fig. 1. The residues, aligning to the conserved aspartate in the motif of PLP-dependent enzymes of fold I, are depicted as stick models. The active site PLP is shown in green, heme in CBS (HEM) is shown in cyan. The scaled up views of the motif within the monomers (left) are accompanied by the motif visualization in the enzyme dimers (right). The following PDB structures are used: serine racemase (SRR) – 6ZSP; serine dehydratase-like (SDSL) – 2RKB; cystathionine β-synthase (CBS) – 4COO. (B) – The sequences corresponding to the motif of dpaL protein, obtained after the structural alignment of the fold II proteins, and the logo corresponding to these sequences. Human L-serine dehydratase/L-threonine deaminase is used to prepare the logo as an additional known PLP-dependent enzyme of fold II (61% identity with SDSL), although its 3D structure is not resolved. Its sequence is added using the Qlustal Omega multiple sequence alignment with the homologous proteins of fold II (B). The residues, aligning to the conserved aspartate in the motif of PLP-dependent enzymes of fold I and in dpaL protein, are shown in bold and marked with an arrow on the motif logo. Used uniprot accession numbers are: P40817 – DpaL; Q9GZT4 – SRR; Q96GA7 – SDSL; P35520 – CBS; P20132 – L-L-serine dehydratase/L-threonine deaminase.

We have also examined a possibility of existence of a potential PNPO-specific motif, using available data on the three PNPO interactors (table). In this case, the score of the found motif is expectedly low due to the small number of the partner proteins. Moreover, the motif is not preserved when the interactor homologs, such as human serine hydroxymethyltransferase, aspartate aminotransferase of *Bacillus sp*. and L-threonine aldolase from *Preudomonas sp*. are added to the sequence search. Last but not least, the motifs thus revealed are not folded into the particular secondary structure, inherent in the identified common motif shown in figs 1 and 2. Thus, the number of known PLP-dependent partners of PNPO (table) is not enough to robustly identify a specific PNPO-interacting site, different from that of PdxK. The same applies to the search of the motif of the PLPBP partners (n=2).

### Pathogenic mutations in tyrosine aminotransferase (fold I) and cystathionine β-synthase (fold II) within or in the vicinity of the identified common motif support its functional significance beyond the active site terminal parts

According to our hypothesis on the relevance of the common motif of the PLP-dependent enzymes for the PLP transfer from its donors, the point mutations within the motif or nearby may be pathogenic, even if they affect the residues outside the active site. Hence, we screened the human variants of the selected PLP-dependent enzymes to find pathogenicity of the human enzyme mutants within or near the characterized motif. For the selection, multiple known mutations of TAT (fold I) and CBS (fold II) have been considered, as dysfunction or dysregulation of these enzymes cause monogenic diseases or malignant transformation, as considered in Introduction.

Of 20 point mutations in TAT causing Richner–Hanhart Syndrome (tyrosinemia type II) [37–43], three are within the motif (A237P) or close to it (L201R, L273P) (fig. 3 A). The characteristic symptoms, such as elevated plasma tyrosine levels and painful hyperkeratotic plaques, occur in patients with L201R and L273P TAT mutations, localized within 5Å from the motif in the alpha-helix and beta-sheet, respectively [41, 43]. Introduction of the positive charge in L201R, or perturbation of secondary structure upon substitution of the beta-sheet leucine residue for proline in L273P, may well prevent the proper protein-protein interaction through the identified common motif, that is shown in purple in fig. 3a. In case of A237P TAT mutation [40], the elevated tyrosine level is accompanied by mild mental disorders instead of skin lesions. Thus, the direct structural impairment of the motif alfa-helix in the A237P mutant is more critical for physiology than the indirect effects of the L201R and L273P mutations in the motif neigbourhood. Remarkably, all the three mutations strongly affect the TAT catalysis despite their positions far away from the active site on the protein surface. This finding is in good accord with presumed functional significance of the identified motif in the PLP transfer.

**Fig. 3.**
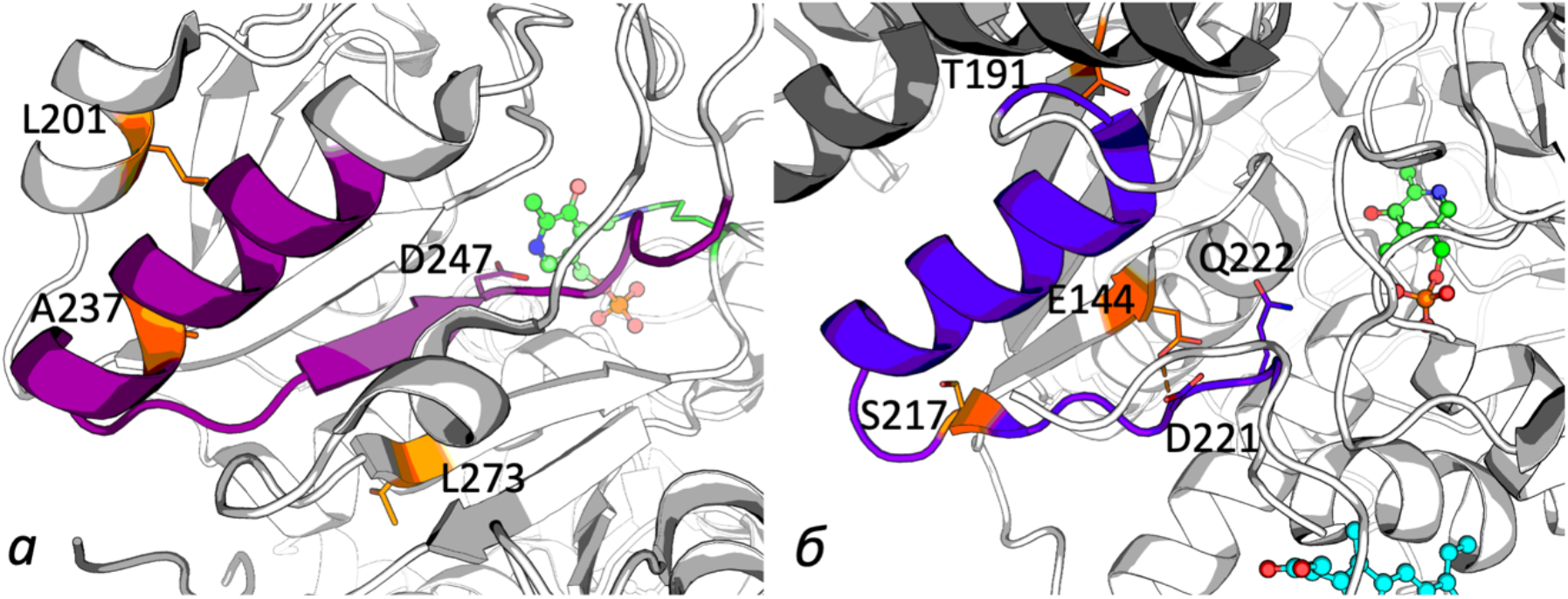
Pathogenic mutations of TAT and CBS within or near the common motif of the PLP-dependent proteins, presumed to be involved in the PLP transfer from the coenzyme donors. (A) The TAT mutations within the motif helix (A237P) and within 5Å from the motif (L201R and L273P). (B) CBS mutations S217F (within the motif), T191M (3.5Å from the motif), and E144K (2.9Å from the motif); H-bonding of E144 to N-atom of the motif D221, preceeding the structurally relevant residue Q222 of the motif (fig. 2 B) in the native CBS is indicated by dashed line.

Of more than 130 point mutations of CBS [21, 44–47], one (S217F) is on the protein surface within the common motif of PLP-dependent enzymes (fig. 3 B). It results in homocystinuria and a 25-fold decrease in CBS activity, compared to wild type [47], supporting the S217 role in formation of the protein complex to deliver PLP from its donors to CBS. Other pathogenic mutations of CBS in the vicinity of the motif and outside the active site are T191M and E144K (fig. 3 B). Their pathogenicity may also be explained by perturbation of the presumed heterologous interaction with the PLP donors and/or PLP transfer process. In case of the mutation T191M this perturbation may be due to the bulkier and more hydrophobic methionine residue, compared to the tyrosine residue in the native CBS. Highly prevalent in Spain, Portugal and South America, the T191M mutation is characterized by varied phenotypic symptoms affecting the skeleton, eyes and CNS [48]. Remarkably, patients with T191M substitution do not respond to the vitamin B6 therapy. The non-responsiveness would be expected for the mutations strongly impairing not the PLP biosynthesis, but the PLP delivery from its donors to the acceptors through the heterologous protein-protein interactions where the identified common motif is presumed to participate. The E144K mutation of CBS changes the charge within a short distance to the motif loop comprising multiple charged residues (fig. 3 B), potentially affecting ionic interactios of the motif involved in the PLP transfer. However, the patients with E144K CBS show positive response to B6 administration. Probably, increased PLP production in the vicinity of the motif, occurring at a higher saturation of the PLP producers with their substrates, may stimulate modification of K144 residue in E144K mutant of CBS by PLP. As a result, the electrostatic perturbations caused by the mutation, may be compensated. The ensuing restoration of the impaired PLP transfer in E144K CBS with the PLP-modified K144 may explain the therapeutic significance of B6 administration in this mutant, absent in the T191M mutant. Another mechanism contributing to the patients’ response to B6 therapy upon mutations affecting the heterologous interactions may be allosteric regulation of the PLP producers by their substrates and products [17, 49], potentially affecting also the heterologous interactions with PLP acceptors.

## DISCUSSION

PLP-dependent enzymes are assigned to seven structural folds by Percudani et al. [1]. 76% of the PLP-dependent enzymes belong to the fold I, followed by the fold II (12%), fold III (5%) and other folds (7%) (fig. 4). The same distribution is inherent in the experimentally confirmed interactions of PLP acceptors with PLP donors (table), where 75% (9 of 12 proteins) belong to the fold I, while the rest 25% are shared between folds II, III and V (8%, i.e. 1 of 12 proteins, for each of the folds). Thus, the confirmed interactions correspond to the incidence of the structural types of PLP-dependent proteins. All the major fold types, including the rare fold V, are shown to interact with the PLP donors, supporting the hypothesis for the PLP transfer in the heterologous acceptor-donor complexes.

**Fig- 4.**
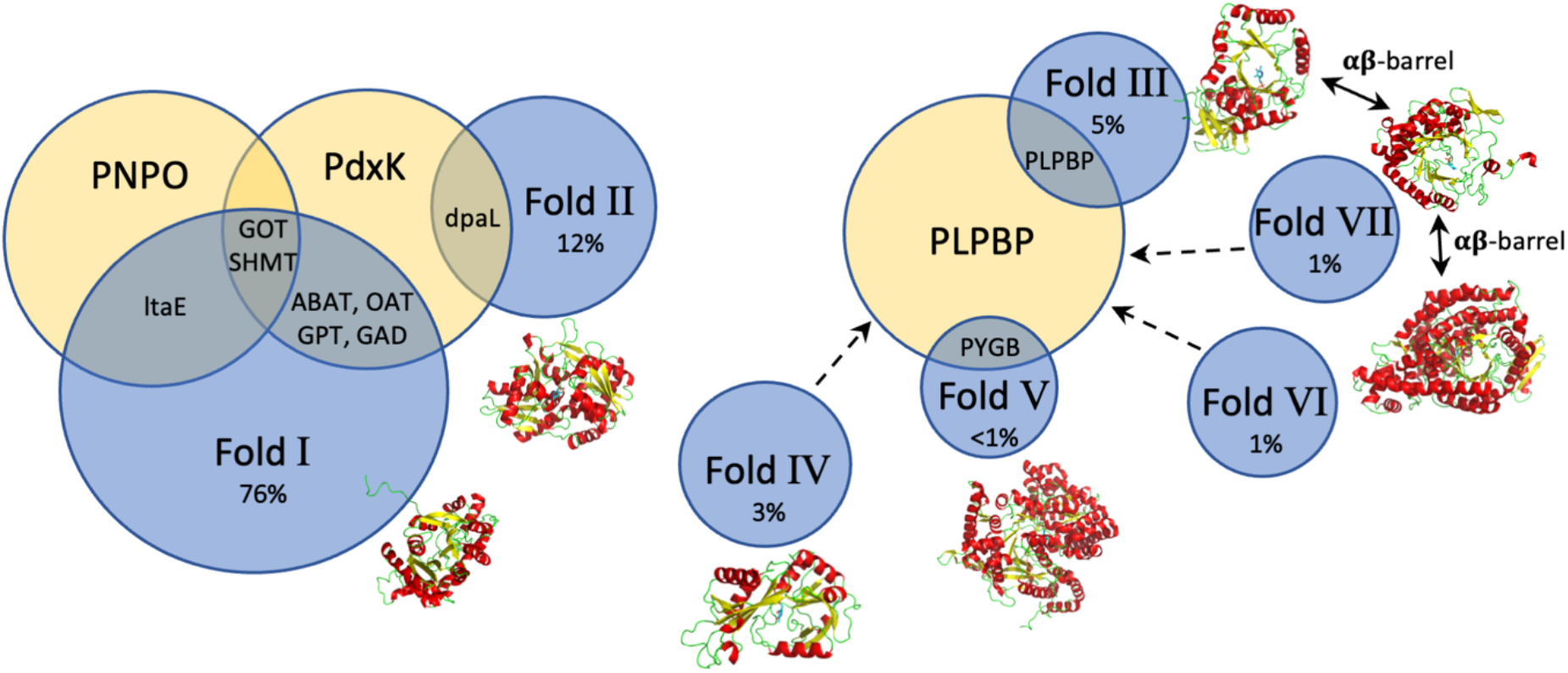
Known (inside the circles) and hypothetical (outside the circles) heterologous complexes between PLP acceptor (blue circles) and donor (yellow circles) proteins. The subunit folds of representative PLP-acceptor proteins are shown near the blue circles. Percentage of the PLP acceptors belonging to each fold, is given according to Percudani et al. [1]). Protein abbreviations: ltaE – L-threonine aldolase, GOT – aspartate aminotransferase, SHMT – serine hydroxymethyltransferase, ABAT – 4-aminobutyrate aminotransferase, OAT – ornithine aminotransferase, GPT – alanine aminotransferase, GAD – glutamate decarboxylase, dpaL – diaminopropionate ammonia-lyase, PYGB – glycogen phosphorylase, PLPBP – PLP-binding protein, PdxK – pyridoxal kinase, PNPO – pyridoxine phosphate oxidase. The suggested interactions of PLPBP with the proteins of folds IV, VI and VII are indicated by dashed arrows.

Existence of the common partners of PdxK and PNPO, i.e. serine hydroxymethyltransferase and aspartate aminotransferase (table), corroborates involvement of the common motif of the PLP acceptors to bind both PdxK and PNPO (fig. 1). However, in view of the limited number of the confirmed protein interactors of PNPO, we cannot exclude that PdxK and PNPO may still have the donor-specific binding sites.

The experimentally confirmed interactions (table) reveal that all partners of PdxK and PNPO belong to the structural folds I and II. Accordingly, the characteristic common motif with specific secondary structure is revealed in these partners of PdxK/PNPO, suitable as an interface for the binding of the PLP donors. In the PLP-dependent enzymes of fold I, the motif incorporates a conserved aspartate residue involved in the PLP binding in the active sites [20]. The corresponding residue is not conserved in the fold II enzymes, where the mode of interaction with the PLP coenzyme is known to differ from that in the fold I enzymes. Nevertheless, also in these enzymes, the C-terminal part of the mptiv is vicinal to the active site PLP, supporting the motif function in the PLP transfer.

In contrast to the PdxK and PNPO interactors, presented by the structural folds I and II, the two known complexes of the third mammalian PLP donor, PLPBP, involve the proteins with the structurally different folds V (glycogen phosphorylase) and III (self-oligomerization) (table, fig. 4). The folds VI and VII, found in a very small number of PLP acceptors [1] and absent in the original classification of the PLP-dependent enzymes [20], share the alpha-beta barrel structure with the fold III (fig. 4). Hence, one may suggest that also these PLP acceptors interact with PLPBP to receive their PLP (shown in fig. 4 by dashed arrows). The fold V is another rare fold of PLP-dependent proteins, shown to interact with PLPBP. Thus, unlike PdxK and PNPO, interacting with the most abundant folds I and II, PLPBP may be required to donate the coenzyme to PLP acceptors of the rare structural folds III-VII (fig. 4).

A recent study of the PLPBP interactors reveals that they are dominated by the proteins participating in the cytoskeleton organization and cell division rather than the PLP-dependent enzymes [33]. The finding suggests that the delivery of PLP to the apoenzymes is not the major PLPBP function. This assumption is supported by the association with the cytoskeleton organization and cell division also of another PLP-binding protein, the PLP-specific phosphatase known as chronophin (PLPP). Apart from hydrolyzing PLP to PL, chronopin acts as a protein dephosphorylase towards cofilin [50], which is a key regulator of actin cytoskeletal dynamics, controlled by the phosphorylation. Association of both PLPBP and PLP phosphatase with the cytoskeleton organization and cell division [33, 50] suggests a currently underestimated role of PLP in these processes, supported by identification of PLPBP orthologs in all kingdoms.

Thus, we have identified a common solvent-exposed protein motif extending to the PLP-binding sites of the majority of the PLP_dependent enzymes. Conformational mobility of the motif upon its binding the PLP donor(s) may be involved in forming the PLP exchange channel. For instance, in glutamate decarboxylase, open conformation of the active site, which has been implicated into the PLP transfer [18], may be stabilized by the heterologous protein complex. Possessing structural similarity in otherwise very different structures of proteins employing PLP (fig. 4), the identified motif may form a universal interface for the protein-protein interactions of the multitude of the PLP acceptor proteins with the limited number of PLP donor proteins. Bringing the active sites of the donor and acceptor proteins close to each other through conformational changes in their heterologous complex, the interaction may enable the PLP transfer between the sites. In case of critical enzymatic functions, the coupled expression of the PLP producers and users may provide an indirect evidence for the complex formation, as the components of the protein complexes associated with fundamental functions, are usually coexpressed [51]. It is worth noting in this regard that in hepatocellular carcinoma, where functions of the PLP-dependent enzymes of fold I (TAT) and fold II (CBS) are decreased [4, 52], also the PLP producer PNPO is downregulated, accompanied by a lower PLP level [53]. Pathogenicity of human mutations outside the active site, but within or nearby this common motif, further corroborates functional significance of the common motif for the PLP transfer to the PLP acceptors.

Although the identified common motif of different enzymes possesses similar structure and certain conserved residues, in each PLP acceptor it also has multiple enzyme-specific residues. These structural features open opportunities for the design of peptide drugs selectively blocking the PLP delivery interface in enzymes of interest, such as CBS, whose upregulation in some cancers is associated with poor prognosis [5, 6].

## CONCLUSIONS

Experimentally confirmed protein-protein interactions and available structural data have enabled identification of a common motif in the multitude of the PLP-dependent enzymes of folds I and II that may provide an interface for their interaction with the PLP-producing PdxK or PNPO. Functional significance of the motif in the PLP transfer between the interacting proteins is supported by pathogenicity of the mutations outside of the active sites, but within or in the vicinity of the motif. PLPBP interactions suggest its binding of the PLP-dependent proteins of the rate structural folds III-VII.

## Contributions

V.A.A. performed literature search, investigation, formal analysis, visualization, and wrote the manuscript draft. V.I.B. supervised the work, analyzed the data, formulated the concept and finalized writing the manuscript. Both authors have read and agreed to the published version of the manuscript.

## Ethics declarations

The study does not include animal experiments or require informed consent statement. The authors declare no conflict of interest. The funding sponsors had no role in the design of the study, as well as in the collection, analyses or interpretation of data, the writing of the manuscript or in the decision to publish the results.

## Electronic supplementary material

The online version contains supplementary material available at https://doi.org...

## Abbreviations

PL: Pyridoxal,
PLP: Pyridoxal-5’-phosphate,
PdxK: pyridoxal kinase,
PNPO: pyridox(am)in-5’-phosphate oxidase,
PLPBP: PLP-binding protein also known as PROSC,
PLP: Pyridoxal-5’-phosphate phosphatase also known as chronophin,
SRR: serine racemase,
CBS: cystathionine β-synthase,
TAT: tyrosine aminotransferase,
GOT: aspartate aminotransferase,
SHMT: serine hydroxymethyl transferase,
ABAT: GABA transaminase,
SDSL: serine dehydrotase-like protein.

