## supplementary figures S1 and S2 for "Protein-protein interfaces as druggable targets: A common motif of the pyridoxal-5’-phosphate-dependent enzymes to receive the coenzyme from its producers"

### Supplementary Materials

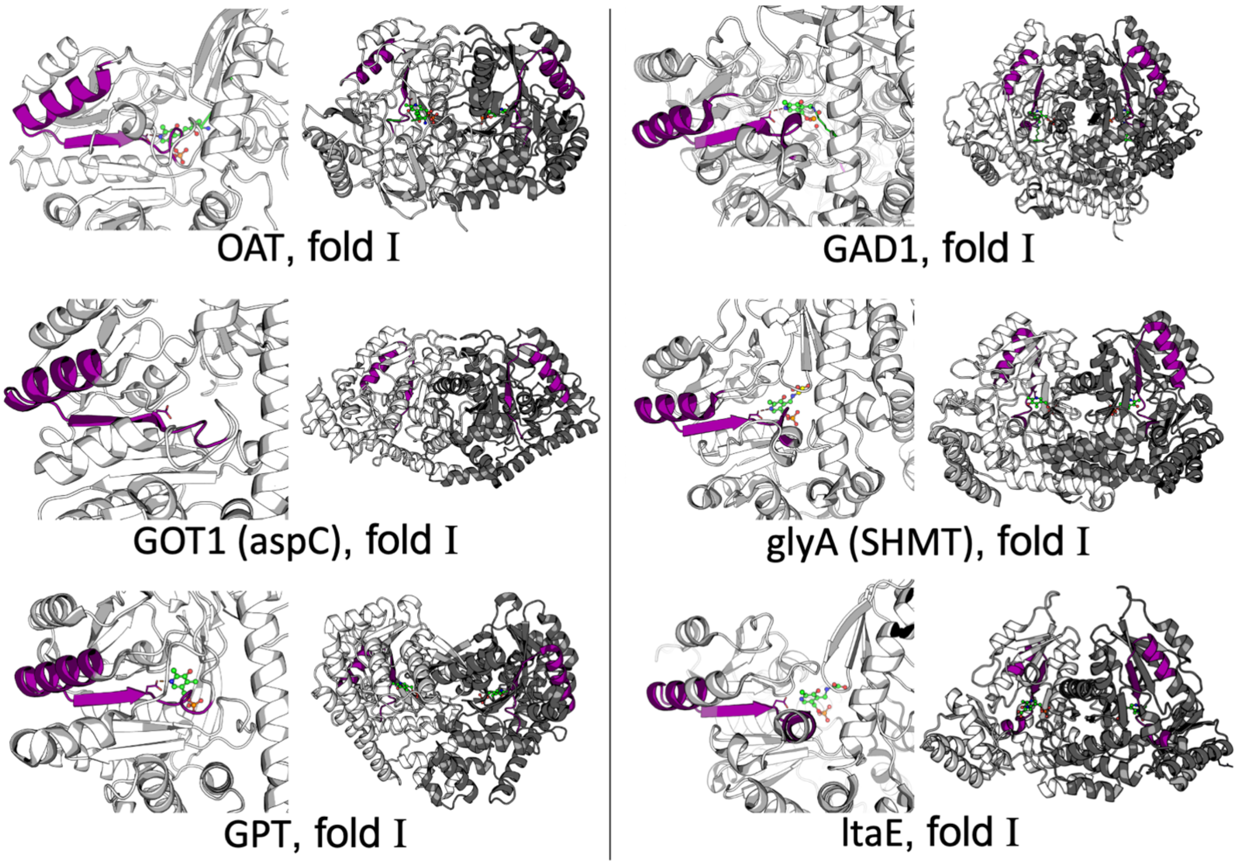


**Supplementary Figure S1.** Localization of the common motif of the known PLK- or PNPO-binding proteins in their 3D structures, additional to the structures of Figure 1. The motif shown in Figure 1 is marked in purple, PLP is shown in green. The localization of the motif in the enzyme dimers (right) is shown near the scaled up view within the monomeric structures (left). For aspartate aminotransferase (GOT) or serine hydroxymethyltransferase (SHMT) one of the two entries present in Table 1 was used. The following PDB structures were used: OAT – 2OAT; GOT1 –3WZF; GPT –3IHJ; GAD1 –2OKJ; glyA – 1KKP; ltaE – 3WLX.

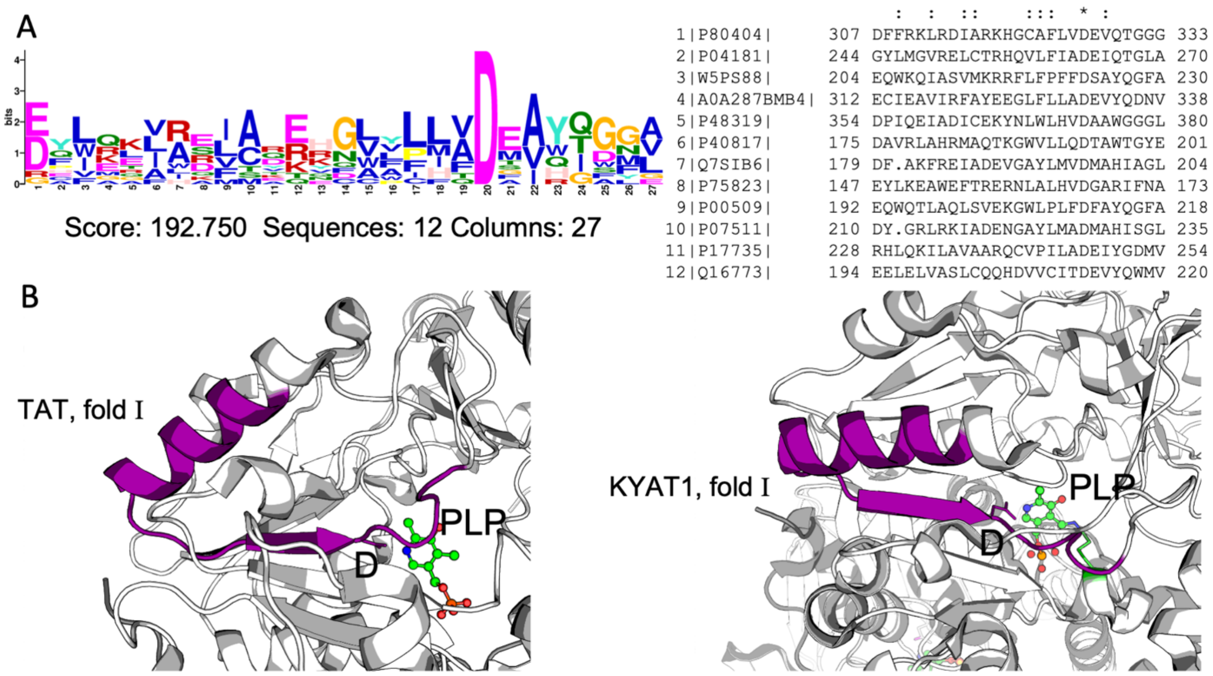


**Supplementary Figure S2.** Identification of the common motif of the known PLP-dependent interactors of PLK and PNPO in the potential interactors belonging to fold I, tyrosine transaminase (P17735, TAT) and kynurenine transaminase 1 (Q16773, KYAT1). A – the logo and the sequences of the motif, found in the PLK-binding mammalian enzymes and the two tested enzymes; B – the localization of the motif in the enzyme, the scaled up view is shown. The following PDB structures were used: TAT – 3DYD; KYAT1 – 3FVS.
